# Extensive fragmentation and re-organization of gene co-expression patterns underlie the progression of Systemic Lupus Erythematosus

**DOI:** 10.1101/2020.01.28.922559

**Authors:** Vasilis F. Ntasis, Nikolaos I. Panousis, Maria G. Tektonidou, Emmanouil T. Dermitzakis, Dimitrios T. Boumpas, George K. Bertsias, Christoforos Nikolaou

## Abstract

Systemic Lupus Erythematosus (SLE) is the prototype of autoimmune diseases, characterized by extensive gene expression perturbations in peripheral blood immune cells. Circumstantial evidence suggests that these perturbations may be due to altered epigenetic profiles and chromatin accessibility but the relationship between transcriptional deregulation and genome organization remains largely unstudied. We developed a genomic approach that leverages patterns of gene coexpression from genome-wide transcriptome profiles in order to identify statistically robust *Domains of Co-ordinated gene Expression* (DCEs). By implementing this method on gene expression data from a large SLE patient cohort, we identify significant disease-associated alterations in gene co-regulation patterns, which also correlate with the SLE activity status. Low disease activity patient genomes are characterized by extensive fragmentation leading to DCEs of smaller size. High disease activity genomes display excessive spatial redistribution of co-expression domains with expanded and newly-appearing (emerged) DCEs. Fragmentation and redistribution of gene coexpression patterns correlate with SLE-implicated biological pathways and clinically relevant endophenotypes such as kidney involvement. Notably, genes lying at the boundaries of split DCEs of low activity genomes are enriched in the interferon and other SLE susceptibility signatures, suggesting the implication of DCE fragmentation at early disease stages. Interrogation of promoter-enhancer interactions from various immune cell subtypes shows that a significant percentage of nested connections are disrupted by a DCE split or depletion in SLE genomes. Collectively, our results underlining an important role for genome organization in shaping gene expression in SLE, could provide valuable insights into disease pathogenesis and the mechanisms underlying disease flares.

**Significance:** Although widespread gene expression changes have been reported in Systemic Lupus Erythematosus (SLE), attempts to link gene deregulation with genome structure have been lacking. Through a computational framework for the segmentation of gene expression data, we reveal extensive fragmentation and reorganization of gene co-regulation domains in SLE, that correlates with disease activity states. Gene co-expression domains pertaining to biological functions implicated in SLE such as the interferon pathway, are being disrupted in patients, while others associated to severe manifestations such as nephritis, emerge in previously uncorrelated regions of the genome. Our results support extensive genome re-organization underlying aberrant gene expression in SLE, which could assist in the early detection of disease flares in patients that are in remission.

**Graphical Abstract:** 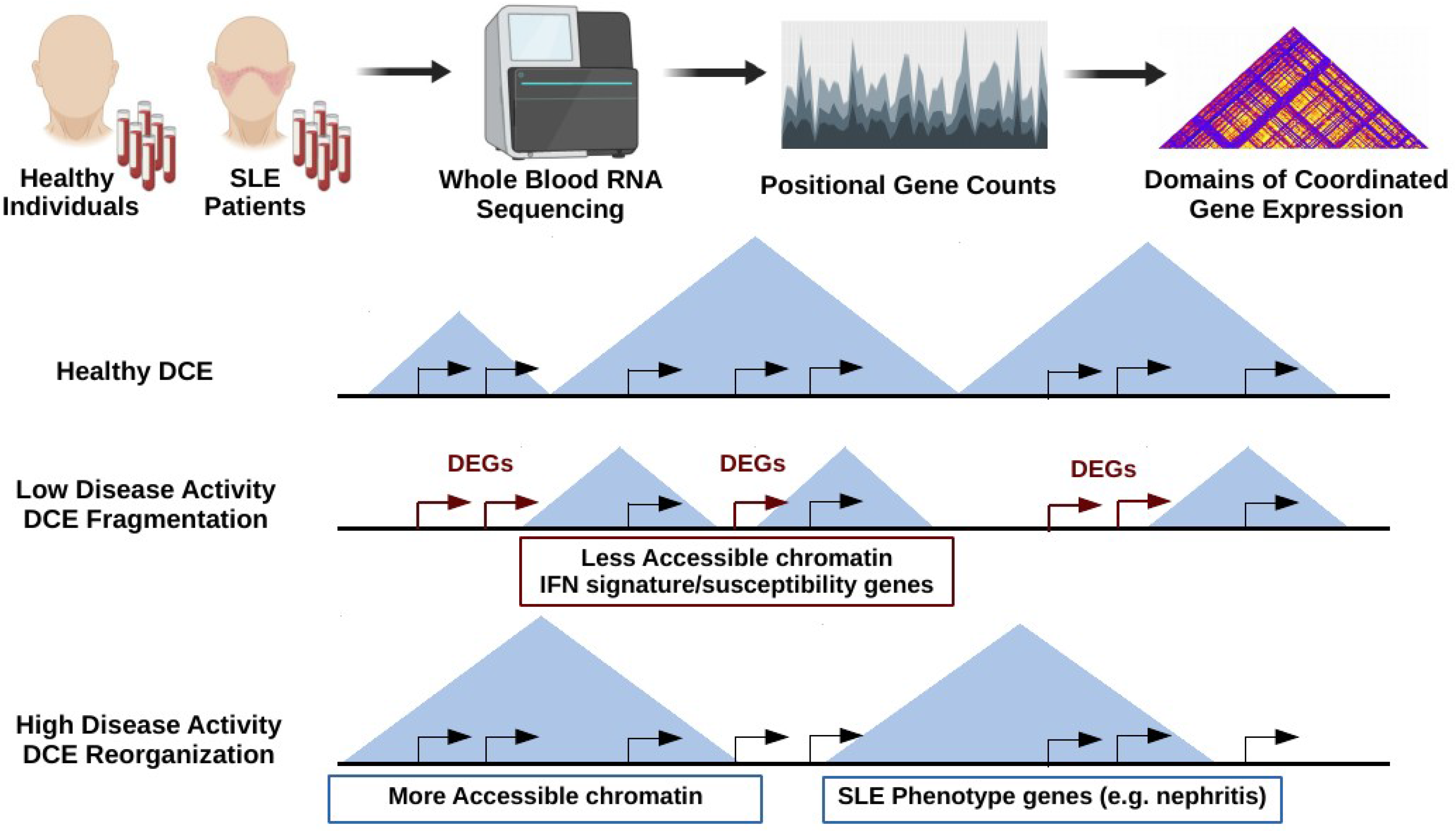

## Introduction

Systemic Lupus Erythematosus (SLE) is considered the prototype of systemic autoimmune diseases due to highly heterogeneous manifestations, variability in symptoms, affected organs and alternating periods of dormancy and increased activity (flares) (1). Several studies of SLE transcription profiles (reviewed in (2, 3)) have reported consistent alterations in key biological pathways, with the Interferon (IFN) signaling pathway being the most prominent example (4, 5). A recent systematic transcriptomic and genetic analysis comparing SLE patients with variable disease activity against healthy individuals, led to the definition of discrete susceptibility and severity gene signatures (6). Beyond gene expression, changes have also been observed at the epigenetic and chromatin levels, with extensive DNA hyper-hydroxymethylation in SLE T-cells (7) and altered chromatin accessibility in naive B-cells from SLE patients under flare status (8).

Given the complexity of the disease at both transcriptome and chromatin levels, an aspect that has not been adequately explored pertains to genome architecture. Over the last years, a number of genomic entities including chromatin loops (9), topologically associated domains (10), enhancer-promoter interacting domains (11), cis-regulatory domains (12, 13) and domains of defined epigenetic characteristics (14, 15) have been shown to define an ever more complex genomic landscape. In spite of their variable size, dynamics and underlying principles governing their creation, a unifying property of these chromosomal entities is the co-ordination of gene expression (16, 17). At the same time, novel high-throughput methodologies have unraveled a strong link between nuclear compartments and transcriptional activity (18, 19). Positional effects in gene expression have been reported since relatively early and their evolutionary and regulation dynamics have been extensively studied (16, 20–22). The importance of gene clustering deregulation in disease has been demonstrated through epigenetics in the case of cancer (23) and genetic associations in the case of Down syndrome (24), but a comprehensive assessment of gene expression clustering has been lacking. Given the apparent extent and impact of genome organization, addressing gene expression changes from an architectural viewpoint could enhance our understanding of the genomic basis of complex pathological conditions, especially those that are accompanied by widespread gene expression alterations, such as SLE.

In this work, we have employed a genomic segmentation approach on an extensive SLE expression dataset (6), aiming to define regions of co-ordinated gene expression for the first time in the context of a complex disease. Our analysis leads to the definition of detailed patterns of transcriptional compartmentalization that vary significantly between SLE and healthy individuals. Interestingly, we find SLE patient genomes to exhibit more fragmented and thus, less structured co-expression patterns, a trend that correlates with the degree of disease activity. The defined *Domains of Co-ordinated Expression* (DCEs) exhibit intricate dynamics, that are associated with both molecular signatures and clinical features of the disease. This represents the first attempt to correlate the complex SLE phenotype with genome topology through detailed transcriptional analysis.

## Results

### Gene co-expression patterns are fragmented in SLE patients

Neighbouring gene expression correlation (25) and modelling (16) have been recently introduced to define how gene expression propagates in space. We employed a topologically-inspired approach to quantify the correlation of gene expression genome-wide. After splitting each chromosome in fixed-size bins, we calculated the transcript count correlation and defined regions of significant co-expression based on a permutation test, followed by local minima localization (see **Methods**). The *Domains of Co-ordinated expression* (DCEs) produced through this analysis are supported by permutation analysis involving 1000 random reshuffling events of transcript counts along each chromosome (see **Methods**, **Figure 1A**). Accordingly, they correspond to statistically robust chromosomal domains, within which gene co-expression is significantly higher as compared to the surrounding regions.

**Figure 1.**
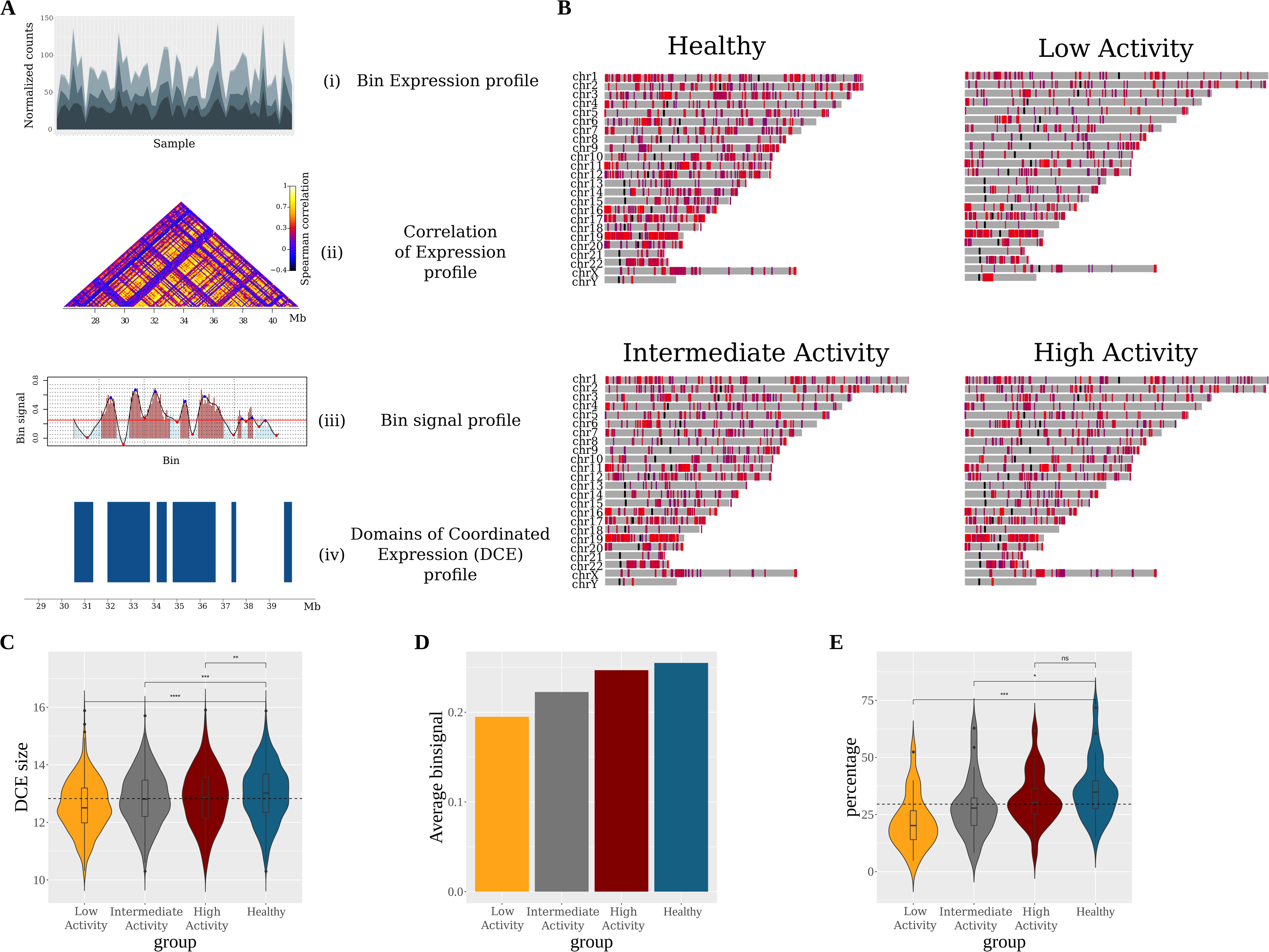
Differential patterns of Domains of Co-ordinated Expression (DCEs) in healthy and patient groups. A. The DCE detection pipeline is represented as a series of ‘transformations’ applied to the expression data. We start by calculating the expression profile of each chromosomal bin using the expression profile of the encompassed genes (i). We then calculate the correlation coefficients between the bins located at the same chromosome (ii). Next, the correlation profile of each chromosome is transformed into a one-dimensional binsignal profile (iii). We analyze that profile, detecting local minima and maxima in order to determine the borders of the domains. Finally, a statistical evaluation of those borders results in the final DCE coordinates (iv). B. Domainograms depicting the distribution of DCEs for the healthy and the three patient groups studied. The color of DCEs represent the respective average binsignal of the chromosomal bins encompassed. C. Violin plots illustrating the estimated distribution of DCE sizes in each group. Classic boxplots are included. The scale of the y axis is logarithmic (log(bps)). D. Average bin signal (co-expression score) for each group. E. Violin plots representing, for each chromosome, the percentage of chromosomal bins that contain genes, with non-zero expression value, and form DCEs. C, E. The results of Mann-Whitney-Wilcoxon tests comparing each patient group to the healthy group are demonstrated by the significance level indicators.

Analysis of DCE patterns between SLE and healthy individuals shows significant differences with SLE gene co-expression being organized into smaller and more fragmented regions. This finding is not confined to specific chromosomes, although gene-dense chromosomes with a more compact transcript pattern show increased overall signal (**Figure 1B**). Notably, DCE patterns correlate with the activity of the disease (SLEDAI). Having stratified patients to three groups according to their SLEDAI values (low, intermediate and high activity) we found that DCE sizes are smaller in low activity patients, where the percentage of the genome organized in DCE does not exceed 9% as compared to 13% and 17%, for intermediate and high activity respectively, and 19% for healthy individuals. Decreased gene co-expression in SLE patients is evidenced by the: a) significantly lower numbers of total DCE for low and intermediate disease activity (**Figure 1B**), b) smaller DCE sizes (**Figure 1C**) c) decreased co-expression signal (**Figure 1D**) and, d) smaller overall percentage of the chromosome covered by DCEs (**Figure 1E**). The observed differences cannot be explained by cell type heterogeneity as shown by an entropy analysis of cell type variability (**Supplementary Figure 1**). The more fragmented expression patterns in low activity SLE genomes are suggestive of generalized perturbations in gene regulation, which could provide mechanistical explanation for the recurrent flares that tend to develop in patients who are inactive. Thus, while a desirable outcome, clinical remission may not necessarily be lacking a molecular fingerprint and the combination of the recently suggested susceptibility signature (6) with our fragmented DCE pattern may provide an interesting framework for the assessment of its stability.

### DCEs are dynamically redistributed in SLE

To gain additional insight into the dynamics of DCEs, we classified DCE patterns into four main groups according to their changes between patient and healthy genomes. We used an implementation of the Jaccard Index to group the domains into: a) DCEs that were left *intact*, b) DCEs that were absent (*depleted*) in patients while present in healthy individuals, c) DCEs that were only present (*emerged*) in patients and, d) DCEs whose coordinates were altered between patient and healthy genomes. The last group was further categorized into DCEs that were *split* (one fragmented into two or more smaller sub-DCEs) or *merged* (two or more joined into one larger) and *expanded* or *contracted*.

Low and high disease activity patients showed the most extensive changes in the pattern of DCEs as compared to the healthy state (**Figure 2A**). A detailed analysis shows that, in agreement with the changes observed at genome-scale level (**Figure 1**), there is extensive fragmentation and redistribution of domains in SLE versus healthy genomes. Contraction and depletion of DCEs are more pronounced in low activity patients, with *contracted* and *depleted* DCEs corresponding to nearly 73% of DCEs in low activity, as compared to 56% and 48% for intermediate and high disease activity genomes, respectively (**Figure 2A**). Conversely, *expanded* and *emerged* DCEs comprise over 30% in high activity versus less than 10% in low activity patients (**Figure 2A**). These observations are suggestive of different modes of dynamic changes in co-expression domains, with low SLE activity genomes characterized by DCE fragmentation and high activity ones featuring a redistribution of co-expression with increased percentages of *expanded* and *emerged* DCEs. This redistribution was also supported by a simple value measure of DCE pattern similarity, calculated with the implementation of BPscore (26), which showed that in spite of being comparable in genome coverage, the DCEs between high activity patients and healthy controls were radically different in terms of genomic localization (**Supplementary Figure 2**).

**Figure 2.**
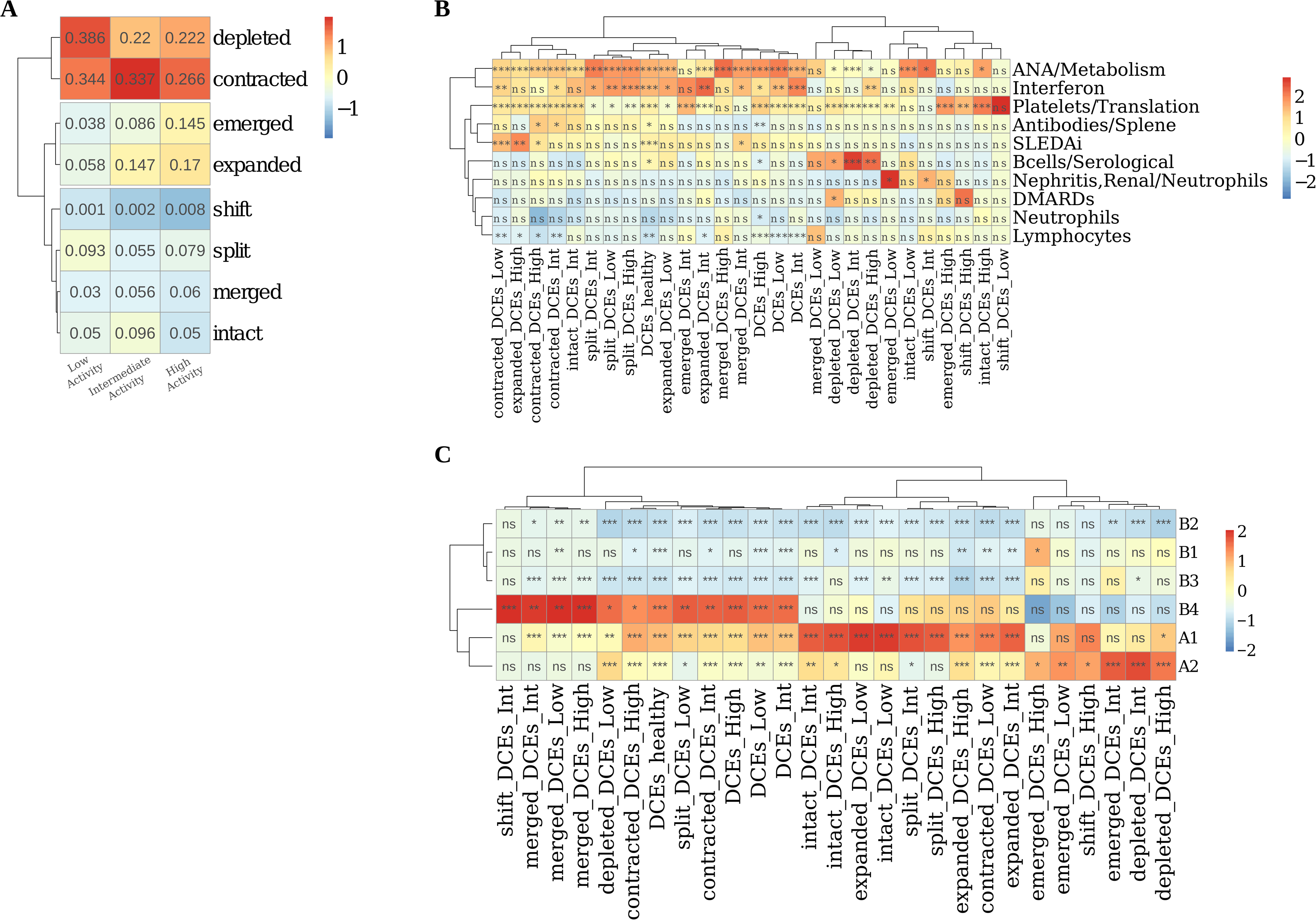
DCEs are extensively fragmented and redistributed in SLE patients and correlate with functional signatures and epigenetic marks. A. Heatmap presenting the different types of DCE reorganization. Numbers inside cells indicate the ratio of the number of DCEs, of the respective type, over the total number of DCEs for each patient group. Colour code is corresponding to column z-score of ratios. B. Heatmap depicting the results of an enrichment test for DCEs in the functionally annotated WGCNA modules. C. Heatmap depicting the results of an enrichment test for DCEs in different genome subcompartments. A-C. Scaling and centering has been performed per column. Trees are illustrating the outcome of hierarchical clustering performed on the data. B, C. Symbols inside cells demonstrate the significance level of the outcome of each test (*:0.05; **:0.01; ***:0.001). Significance has been assessed by a non-parametric, permutation-based test.

### Gene expression changes are reflected upon DCE dynamics

Changes in the patterns of co-expression may be linked to differential gene expression and underlying chromatin dynamics. To address this, we employed Modular and Weighted Gene Co-Expression Network Analysis (WGCNA) (27) of differential gene expression on the three disease activity groups against healthy individuals. The results were strongly suggestive of quantifiable phenotypic variability between patients with different clinical activity states, in agreement with the previously defined susceptibility and severity gene signatures (**Supplementary Figure 3**). In addition we were able to define gene expression modules and to correlate them to clinical characteristics such as disease activity. Comparison of WGCNA results with clinical characteristics of the cohort samples allowed us to identifiy a “*SLEDAI gene module*”, which comprised 224 genes, enriched for innate and adaptive immune pathways, particularly signaling through the Fc-γ and B-cell receptors (BCR). A “*Nephritis module*” (184 genes) and an “*IFN module*” (282 genes) were also identified, the latter being highly associated with anti-nuclear and anti-DNA antibodies (**Supplementary Figure 4**).

In order to investigate how changes in the patterns of co-expression may be linked to differential gene expression we analyzed the degree of overlap between the different DCE categories and the WGCNA modules obtained from our dataset (**Figure 2B**). The *IFN module* was over-represented in *split* DCEs across all SLE groups and was particularly enriched in DCEs that become *depleted* within the high activity group, implying that increased IFN pathway gene activity may be linked to loss of co-expression structure. Conversely, the *nephritis/neutrophil-specific module* was enriched in *emergent* DCEs from low disease activity genomes, which is suggestive of gene deregulation being also associated with the creation of new co-expression domains. The correlation of *nephritis/neutrophil-specific module* with emergent domains in low activity genomes could be indicative of underlying tendencies in gene deregulation present even in patients without developed symptoms who may yet be predisposed to flare. Consistent with observations at the level of functional enrichments, the *B-cell module* was enriched in DCEs that are depleted being largely absent from high disease activity DCEs. Taken together, these findings indicate that functional aspects of gene expression pertaining to distinct clinical characteristics are reflected on the genome organization.

### DCE dynamics are strongly associated with chromatin accessibility and chromosomal compartments

Transcriptional coordination in self-contained domains is tightly linked to underlying chromatin organization at various levels ranging from topologically associated domains (TADs) to more extended chromomomal compartments. We thus went on to correlate the dynamics of DCE patterns with underlying genomic features related to chromatin accessibility and chromosomal compartments. By comparing the coordinates of stable and dynamic DCEs against ATAC-Seq peaks defined for B-cells in severe-case SLE against healthy individuals (8), we found *split*, *contracted* and *merged* DCEs (of all disease activity groups) to be enriched in peaks of decreased chromatin accessibility (**Supplementary Figure 5**). Conversely, *depleted* and *emerged* DCEs of all SLE activity groups were enriched (although with a smaller effect size), almost exclusively, in over-accessible regions. This finding signifies a clear distinction between the DCEs that are locally modified, which tend to be confined in under-accessible regions, and those that are dynamically re-distributed, which are preferentially located in more accessible chromatin.

We performed a similar analysis at the level of chromosome compartments (at 100kbp resolution) as defined in a B-lymphoblastoid cell line (9). On a large scale, chromosomes may be organised into two broad compartments labelled A and B, corresponding to active and inactive chromatin, and also bearing other distinct properties. These may be further subdivided to A1 and A2 and B1 to B4 (9). A chromosomal coordinate overlap enrichment analysis showed DCEs to be generally enriched in the euchromatic A compartment (**Figure 2C**). When focusing on specific DCE subtypes, we found that regions belonging to the most dynamic subsets of *emerged* and *depleted* DCEs were enriched in the A2 subcompartment, which is associated with late-replicating, low GC content DNA, enriched in H3K9me3 and longer gene transcripts (9). On the other hand, intact DCEs and in general, DCEs that are less dynamic appear to be more enriched in the gene dense, early-replicating A1 subcompartment. Enrichments in the B4 subcompartments are probably due to the over-representation of particular DCEs in chromosome 19, which hosts the entirety of this very small subcompartment.

Together, the differential enrichments of *split* and *contracted* DCEs, compared to the dynamically redistributed *emerged* and *depleted* regions, in terms of chromatin accessibility and genome compartments, indicate an interplay between gene regulation and underlying chromatin environment. Regions of high gene density tend to have highly correlated gene expression, but that this pattern changes radically with the splitting of co-expression domains in low disease activity and the emergence of new, probably re-arranged domains in high disease activity SLE patients. We hypothesize that epigenetic changes that increase chromatin accessibility, in particular in A2 genomic compartments, may create a permissive environment for the redistribution of co-regulated genomic domains, which are, moreover, associated with functions characteristic of increased disease activity.

### DCE splits are driven by differential expression of transcriptional regulators and disrupt enhancer-promoter interactions of key biological functions

While *split* DCEs represent no more than 5-10% of the total genome coverage, they are highly enriched among differentially expressed genes and in particular with the *IFN module*. Given their additional enrichment in low disease activity patients and therefore, their possible implication in further disease progression, we performed a focused analysis of *split* DCEs and the genes lying on their boundaries (see **Supplementary Methods**). The latter were predominantly enriched among the targets of specific transcriptional regulators, a number of which are associated with zinc finger factors (SALL1, Ikaros, ZIC3 etc.) and oncogenes (GLI1, ING4) (**Figure 3A**). Members of the Ikaros transcriptional regulators have been genetically associated with SLE (2), and interestingly, *IKZF3* lies within a disrupted DCE in all SLE groups.

**Figure 3.**
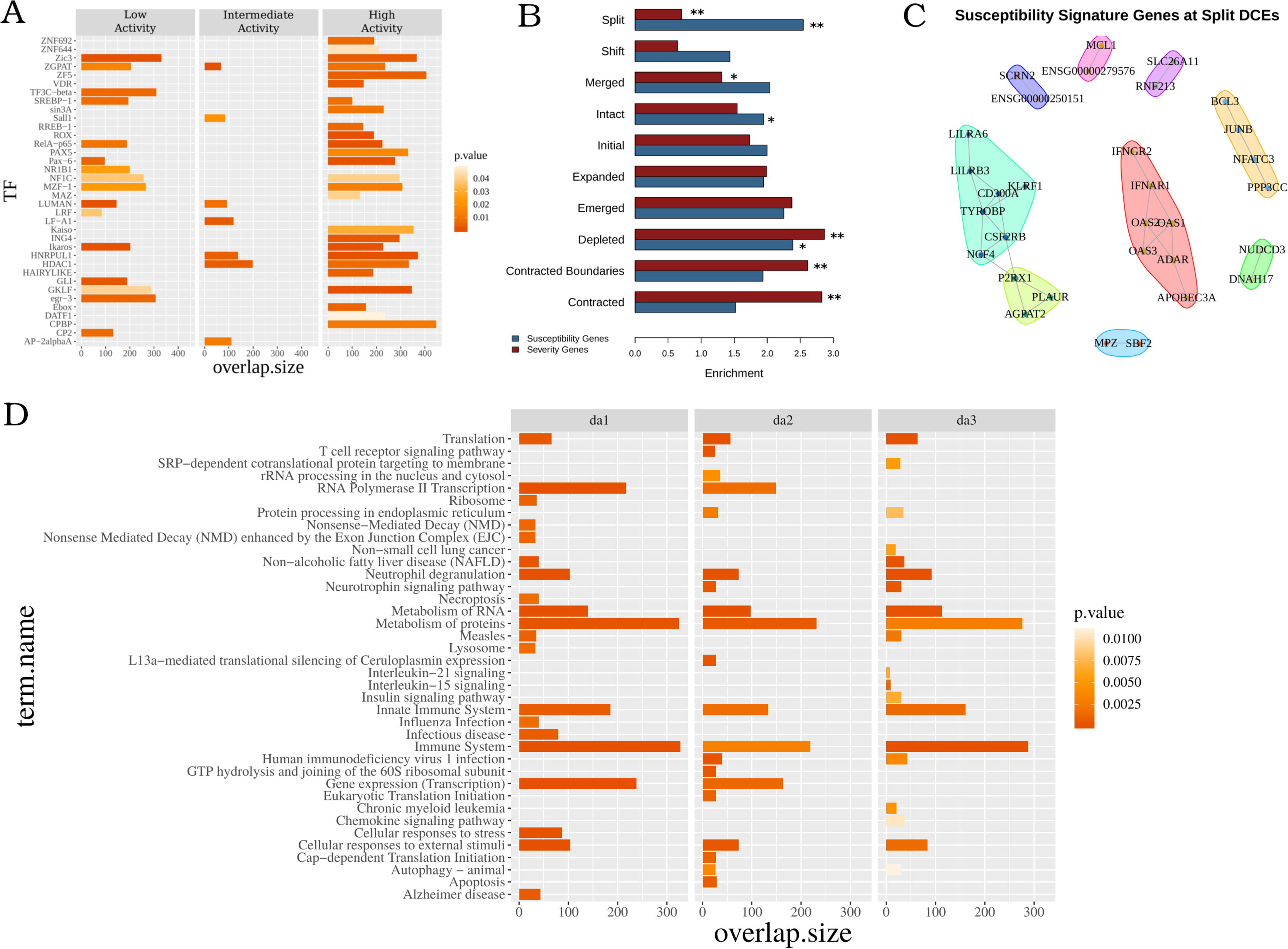
Functional analysis of the disruption events. A. Enrichment analysis of ‘Disruptors’ in genes that are commonly regulated (suggested by the mutual regulatory motif matches - TRANSFAC database) by transcription factors indicated on y axis. The overlap between the query gene set and the corresponding Pathway members or TF-target genes are displayed on the x axis. The color of each bar illustrates the corrected p-value of the corresponding enrichment test. B. Average positional enrichments of susceptibility and severity genes from (6) against different types of DCEs. Significance levels of one hundred permutations (*:0.05; **:0.01). C. Protein interaction networks for susceptibility signature genes that are found to be differentially expressed and overlapping split DCE boundaries, as obtained from STRING-DB (28). Genes are grouped on the basis of a modularity analysis. Modules are shown with coloured polygons around genes (red: interferon signature genes, cyan: DAP12 signaling, lime: neutrophil module, green: B-cell module) D. Pathway enrichment analysis of genes which correspond to enhancer-tss links (CD4+ cells - Enhancer Atlas), that are nested in healthy group DCEs but disrupted in SLE. The top 20 most significant KEGG or/and REACTOME pathways are presented.

Based on the differences in the extent of split DCEs between low and high activity genomes, we next assessed their overlaps with the SLE susceptibility and severity gene signatures as previously defined (6) for the same dataset. We found significant differences between the two gene sets with *susceptibility genes* being highly enriched in *split* DCEs in contrast to a depletion of severity genes (**Figure 3B**). Genes belonging to the *susceptibility signature* are also enriched in the subset of differentially expressed genes that are found in low disease activity *split* DCE boundaries (p=0.0061). Protein and regulatory interaction network analysis of these genes, performed through STRING-DB (28), revealed an IFN gene signature (**Figure 3C**) and interestingly, a set of highly connected genes associated with DAP12 signaling (**Figure 3C,** cyan polygon). DAP12 (TYROBP) is a key activator of NK cells, which are reported to have impaired function in SLE patients (29). Smaller network modules were associated with neutrophils (lime) and B-cells (yellow). We may thus see how, by focusing on split DCE regions we may prioritize genes of the broader susceptibility signature and to investigate their functional connections.

Given the DCE definition as regions with increased regulatory interactions, it is plausible to expect that gene promoters are more likely to be associated with enhancers that are lying within the same region. To test this hypothesis, we obtained cell-type specific promoter-enhancer interactions for CD4, CD8 and CD14 and CD19 cells from Enhancer Atlas (30) and identified genes whose promoter-enhancer pairs were nested within the same DCE in the healthy state but disrupted in SLE. We found that a significant percentage of enhancers-promoter connections that are completely nested in healthy DCEs are disrupted by a DCE *split* or *depletion* in one of the SLE disease activity states. Thus, it seems that the redistribution of gene co-regulation domains in disease may also be disrupting the regulatory links between enhancers and their cognate promoters. Functional enrichment of the genes, whose enhancer-promoter associations are disrupted in SLE revealed relevant biological functions (**Figure 3D**). More specifically, functions related to the *immune system* are, as expected, highly enriched in all disease activity groups. Others, such as *protein metabolism*, *translation* and *protein turnover* are particularly enriched in high disease activity patients. *Gene transcription* is enriched in low and intermediate activity but absent from high activity genomes, suggesting major changes in general functions. Interestingly, interleukin-16 (*Il15*) and interleukin-21 (*Il21*) signaling are specifically enriched in high activity patients even though with low effect sizes (**Figure 3D**).

## Methods

### Definition of domains of coordinated expression (DCEs)

To call domains of co-ordinated expression, we modified a methodology that was introduced for the definition of topologically associated domains (TAD) (31), in our case, by using expression correlation data instead of chromosomal contact frequencies. Statistically robust expression correlation matrices (see **Supplementary Methods**) were used as input. Domains of Co-ordinated Expression (DCEs) were defined as genomic regions of consecutive chromosomal bins with correlation above average, delimited by statistically significant boundaries. More specifically, DCE detection is a four-step pipeline, which is repeated for each chromosome and for every study group (see **Figure 1A**).

1. First, we compute a signal that runs along the chromosomes and is indicative of the local average correlation of expression. We achieve this by sliding two juxtaposed windows of equal size along a chromosome with a single-bin displacement, until the whole chromosome has been covered. In every iteration, we use the correlation matrix that has already been constructed and statistically evaluated. We look up the correlation values concerning the relationship of the two regions and calculate their average. That value is assigned to the chromosomal bin located in the middle, more precisely, the downstream-most bin inside the upstream window.
2. Subsequently, the calculated signal is used to detect DCE boundaries. Hence, the second step of the pipeline is to compute a smoothed function of that signal, using a smoothing spline, and to detect all local minima of that function. DCEs are initially detected as regions between local minima with a value lower than 0.25, which is the average genome signal for the healthy, control group.
3. The third step is to statistically evaluate and refine the boundaries. We estimate the significance of the boundaries by utilizing a Mann–Whitney U test to compare “within” and “in-between” correlation coefficients. In case any of the initially calculated boundaries does not reach the required statistical significance threshold (p-value > 0.05), we “chop” that boundary by one bin towards the centre of DCE, and repeat the test. DCEs with any remaining non-significant boundary are discarded.
4. The last step includes the fusion of neighbouring DCEs which are separated by up to two bins with signal lower than 0.25. The maximum number of bins, with a signal lower than 0.25, allowed in the final DCE is two, thus allowing at most two such fusion events. This step enhances the robustness of the pipeline and decreases the noise in our data. The window size used in bin-signal calculation and in boundary evaluation was three bins, based on the maximization of average intra-DCE correlation of chromosome 1 of the healthy group.

## Discussion

Genome organization is intricately linked to gene expression and regulation in health and disease, with differentially expressed genes creating clusters under various conditions. Our study, the first such conducted in SLE, shows that genes are organized in extended domains of coordinated expression but, moreover, that these domains are highly dynamic and extensively reorganized during disease progression. While high activity patient patterns are suggestive of a general re-organization of gene regulation that extends to broader chromosomal domains, increased fragmentation of gene co-expression is observed even in the genomes of patients with very low disease activity. This may suggest that the observed disruptive patterns of gene expression may be related to the way initial cellular signals propagate in the genome in order to affect hundreds of abnormally regulated genes. Thus, the more disconnected co-expression in low activity SLE genomes could be linked to mechanisms, with which flares occur even in patients that are in remission.

While, the governing principles of such mechanisms are yet to be resolved, our analyses suggest a key role for the chromatin environment. Differential enrichment of DCE patterns between open and closed chromatin and chromosomal compartments pertaining to early and late-replicating chromatin, are strong indications of epigenetic patterns underlying the fragmentation and re-organization of gene co-expression. Epigenetic effects, downstream of environmental triggers are expected to lie at the basis of SLE aetiopathogenesis, given the limited association of genetic factors reported for the disease. Further investigation of the mechanisms linking chromatin structure and the organization of gene expression in SLE could be assisted by our approach, through the prioritization of chromosomal domains with increased regulatory potential.

Besides epigenetic phenomena, the formation of co-expression domains could occur more transiently as the result of differential expression in any given setting (32), through the clustering of differentially expressed genes, that have been spatially constrained through evolution (20, 33). Such a notion is supported by our data in two ways. First through the association of the observed DCEs with functions that are known to be activated in SLE. Major pathways related to the intensity of the symptoms (such as the IFN signature) are associated with the disruption of co-expression, while downstream effects of SLE, related to the damage of organs (e.g. nephritis) are correlated with the general re-organization of co-expression in emergent domains. In addition gene signatures from both expression (6) and genome-wide association data are enriched in various types of DCEs (**Supplementary Figure 6**), a strong indication that transcriptomic as well as genetic data may reveal a hidden layer of information when studied through the lens of genome organization.

The dynamics of co-expression clustering are also linked to differential expression, through the tendency of deregulated genes to occur in the boundaries of split DCEs. Inspection of the dynamics of DCE *splits* is, moreover, indicative of the general pattern of fragmentation and redistribution as is showcased in a number of examples where, compared to a contiguous DCE pattern in the healthy state, we observe splits in low disease activity and more generalized reorganization in high disease activity patients (**Figure 4**). The fact that split/disrupted regions are more prominent at low disease activity genomes, combined with their proximity to genes belonging to the susceptibility signature, may come as an indication of an underlying hierarchy behind the gene deregulation program. Indeed, we find enhancer-promoter associations of high relevance to be possibly affected by the disrupted patterns of gene co-expression, which is strongly indicative of DCE splits having a possible multiplicative effect on gene regulation.

**Figure 4.**
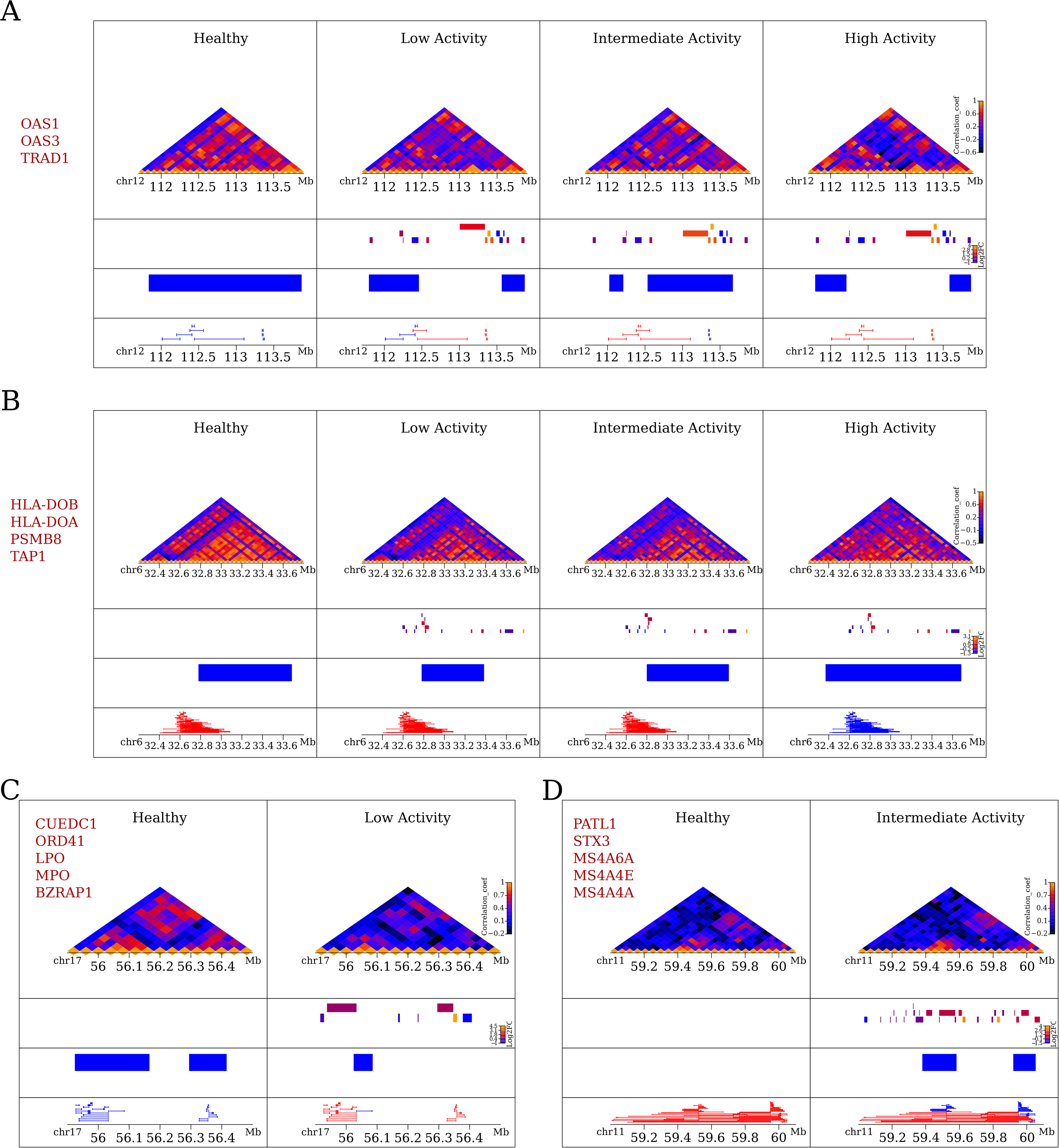
Examples of alterations in the co-expression profile. Heatmaps of expression correlation for selected loci for characteristic cases of disrupted (top), expanded (middle), deleted (bottom left) and emerged DCEs (bottom right). Heatmaps were created with the Sushi package from Bioconductor. Values in heatmaps correspond bin signal, while the tracks below them show (from top to bottom) gene positions colour-coded for differential expression as log2(fold-change), DCE coordinates and enhancer-promoter associations that are entirely included in the same DCE (in blue) or not (in red). Names of differentially expressed genes in each locus are shown on the side of each panel.

The approach we present here constitutes a first attempt to analyze gene expression at the level of genome organization in a complex disease and points to a number of interesting hypotheses linking the SLE phenotype with the underlying genome structure. Targeted conformation capture experiments on homogeneous cell cultures could be implemented in order to test these hypotheses. At the same time, the implementation of single-cell approaches at both transcriptome and genome conformation levels, could provide a data-rich framework for the application of our approach, with the final aim of obtaining cell-type specific co-expression profiles at increased resolution.

## Data Availability

Original RNASeq data have been deposited at the European Genome-Phenome Archive (EGA) under the accession number EGAS00001003662. Processed data and original code for all presented analyses may be found at https://github.com/vntasis/SLE_spatial_gene_expression

## Acknowledgements

We would like to acknowledge the contribution of Halit Ongen, Irini Gergianaki, Maria Trachana, Luciana Romano-Palumbo, Deborah Bielser, Cedric Howald, Cristina Pamfil, Antonis Fanouriakis, Argyro Repa and Prodromos Sidiropoulos in patient screening/enrollment and data generation.

