## Supplementary Information for "Extensive fragmentation and re-organization of gene co-expression patterns underlie the progression of Systemic Lupus Erythematosus"

### Results

#### DCE patterns are not related to cell type heterogeneity

One thing that needs to be stressed is that our approach is based on gene expression from a heterogeneous cell population. In order to assess the extent, to which the differences in the DCE patterns and the more fragmented distribution in SLE patients may be attributed to cell population heterogeneity we used the inferred cell type distribution acquired by (1) for the same dataset. We actually found healthy samples to be more heterogeneous in terms of different cell type representations (**Supplementary Figure 1**). That is, the more fragmented patterns stem from the less homogeneous cell samples and therefore cannot be attributed to cell-type expression variability.

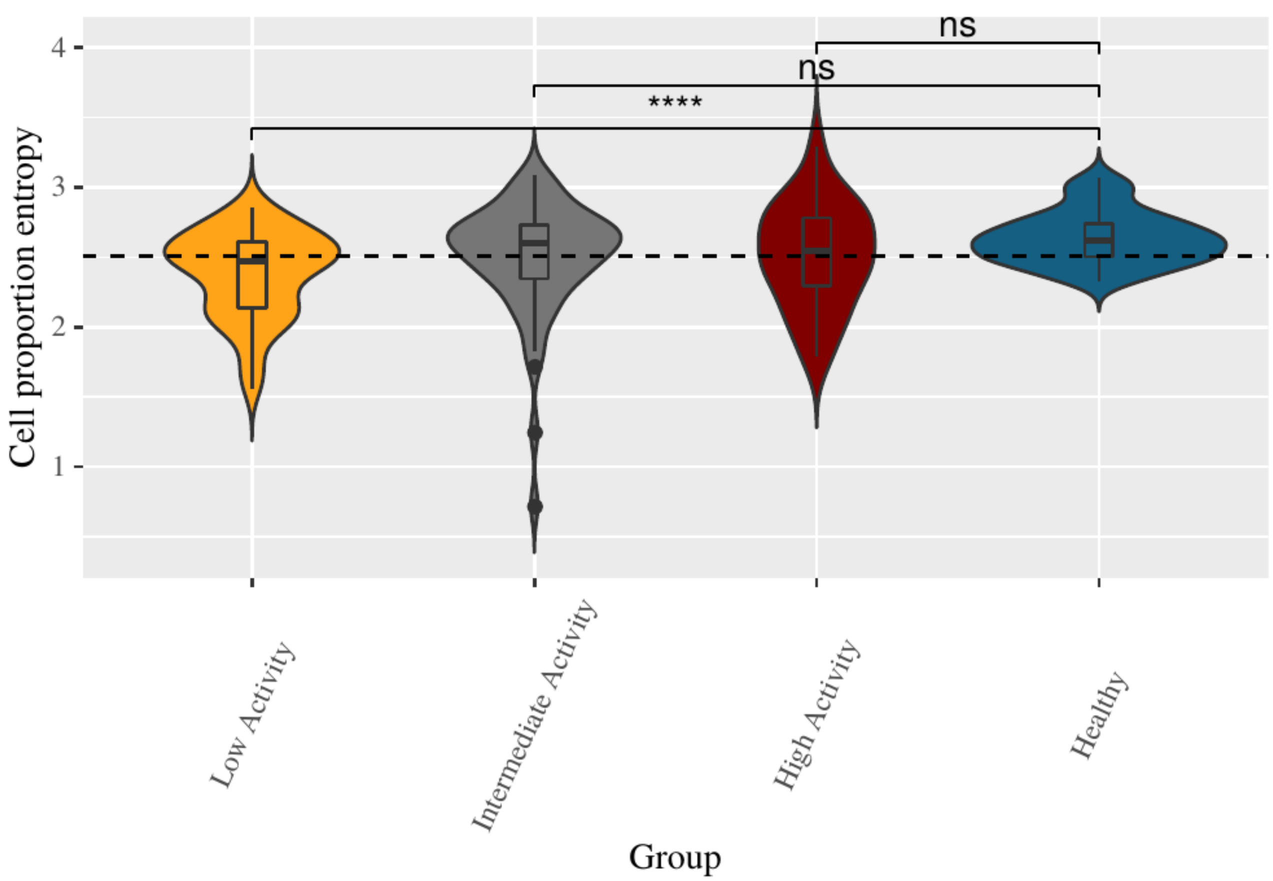


**Supplementary Figure 1. Cell proportion distribution.** Violin plots presenting the estimated distribution of cell proportion entropy (see Methods section) per group studied. The results of Mann-Whitney-Wilcoxon tests comparing each patient group to the healthy group are demonstrated by the significance level indicators. Classic boxplots are included. Dashed line represents the overall average value of the depicted variable.

**BP-Score comparison**

A simple value measure of DCE pattern similarity may be obtained with an assessment of genomic coordinate changes through the implementation of BPscore (6), which allowed us to see that even if the high-activity DCEs are comparable in terms of genome coverage with the healthy ones, they were radically different in terms of coordinates as suggested by the BP scores (**Supplementary Figure 2**). High BP scores are also observed when comparing high with low activity genomes (data not shown) a fact that is strongly suggestive of a general re-distribution of co-expression in different domains, accompanying the more extensive gene expression changes in high activity patients.

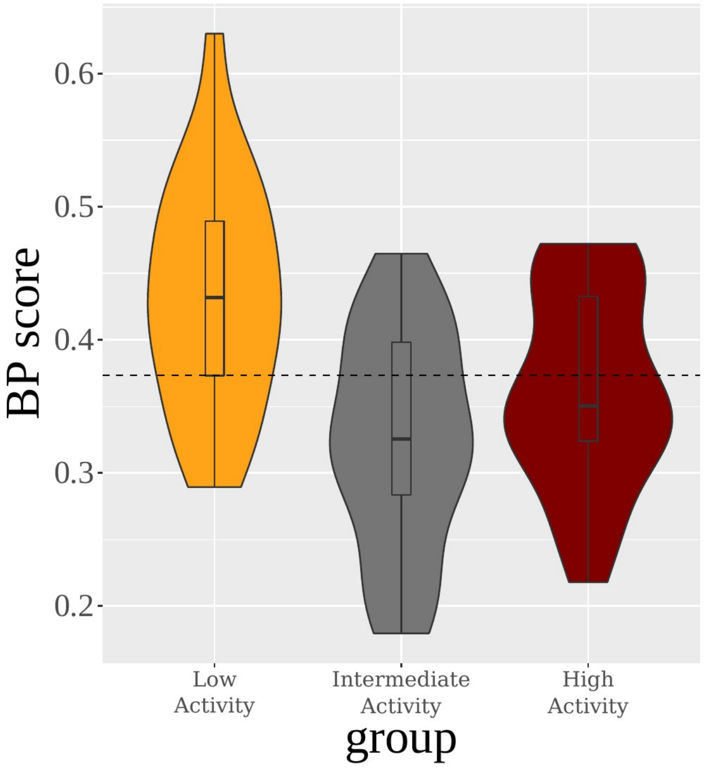


**Supplementary Figure 2 BP distance score characterizing the different patient groups.** Violin plots presenting the estimated distribution of BP distance score (per chromosome) between the DCE profiles of patient groups and the healthy control group. Classic boxplots are included. Dashed line represents the overall mean BP score value.

**Modular Anaysis of Differential Gene Expression and Weighted Gene Co-expression Network Analysis (WGCNA) identify gene sets that are associated with distinct clinical manifestations of SLE**

We used a recently published gene expression dataset on SLE (1) to assess the differential expression levels (against healthy individuals) for three distinct patient groups according to disease activity based on the SLEDAI index (see **Supplementary Methods**). We employed a modular analysis of differential gene expression (**Supplementary Methods**) which uncovered quantitative differences in key pathways across disease groups. Functional modules strongly associated with SLE, such as interferon signaling, neutrophil activation and the innate antiviral response, showed gradually increasing enrichments from low to high activity state (**Supplementary Figure 2**). More general biological functions such as cell cycle, primarily associated with T-cell division, become enriched only in high activity patients. Functions pertaining to plasma cells and B-cells were enriched in under-expressed genes, yet this enrichment was inversely associated with disease activity (**Supplementary Figure 3**. These results are suggestive of quantifiable phenotypic variability between patients with different clinical activity states, in agreement with the previously defined susceptibility and severity gene signatures (1).

We took advantage of the detailed clinical information (including hematological and immunological data and macroscopic observations) that was available, by combining it with a *Weighted Gene Co-expression Network Analysis* (WGCNA, see **Supplementary Methods**). We identified significant co-expression modules, that is, groups of genes that tend to have similar expression levels across healthy individuals and SLE patients. These modules were then associated with available clinical traits at single-patient level, in order to define gene sub-signatures of clinical relevance. A number of gene modules were highly correlated with disease activity (SLEDAI), while distinct gene sets correlated with clinical traits such as active nephritis, biomarkers such as anti-double stranded DNA antibodies, cell-specific attributes for neutrophil, plasma and B-cells, as well as molecular pathways such as the interferon signaling (**Supplementary Figure 4**).

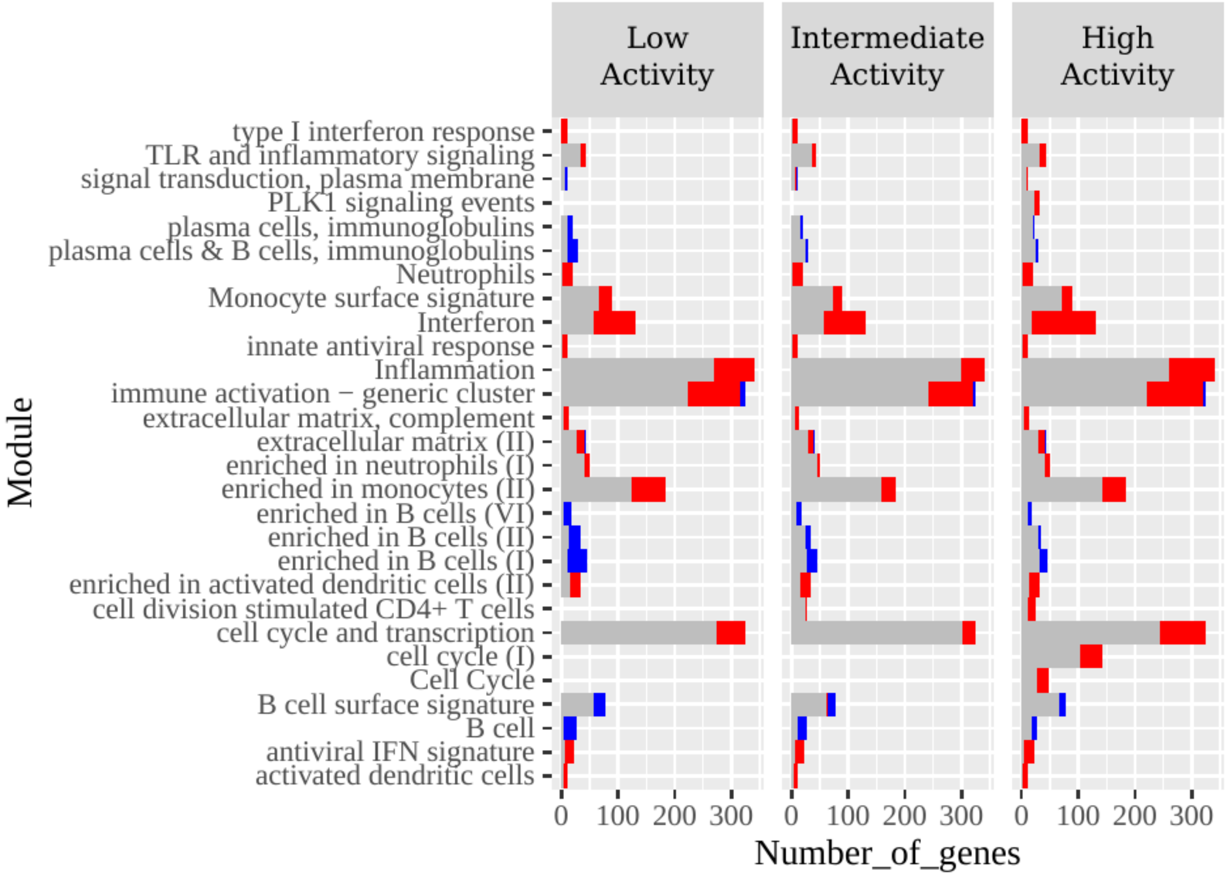


**Supplementary Figure 2**. **Gene set enrichment analysis of differential patient expression.** Barplots illustrating the results of gene set enrichment for blood modules (see Methods). Genes expressed in each patient group were sorted according to their log-FoldChange (calculated in regard to the control healthy group) for this analysis. Top significant modules for each patient group, based on the CERNO test (corrected p-value <= 0.05) (8) are presented. The overall length of each bar illustrates the total number of genes, which are members of the respective module. Overexpressed DEGs are represented by the red part on each bar. Underexpressed DEGs are represented by the blue part on each bar.

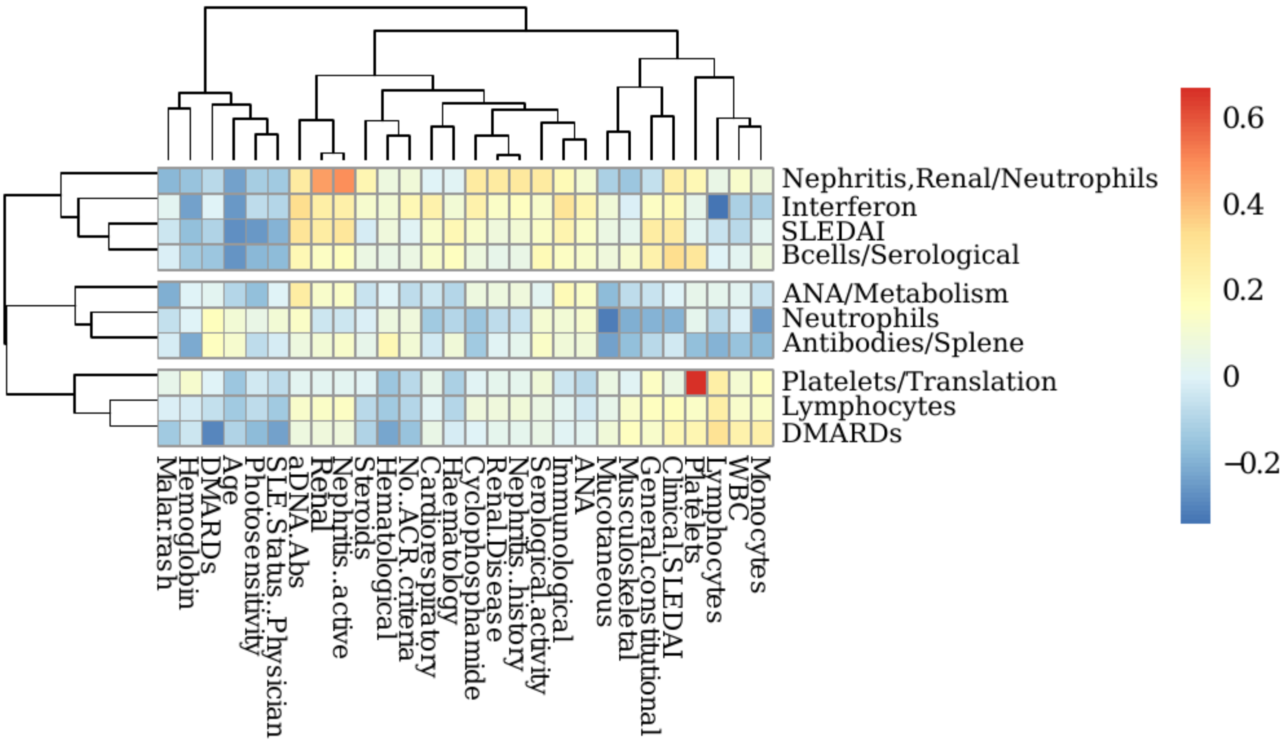


**Supplementary Figure 3. Correlation of WGCNA modules with different clinical traits.** Heatmap depicting correlation values calculated between a variety of patient clinical traits and modules identified by their WGCNA eigengene vectors. Modules are represented by the rows of this matrix and have been named according to their most significant correlated trait and/or a pathway enrichment analysis. Trees are illustrating the outcome of hierarchical clustering performed on the data.

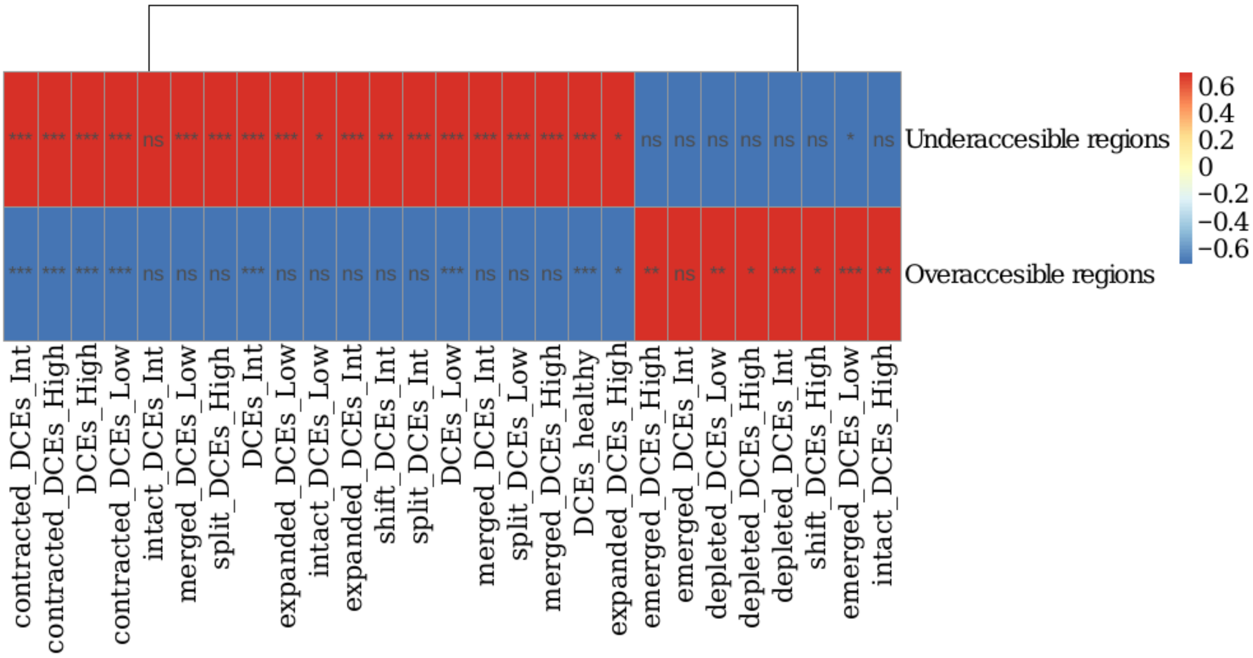


**Supplementary Figure 5. Enrichment of DCEs in differentially accessible regions.** Heatmap illustrating the results of an enrichment test for DCEs in differentially accessible regions. Scaling and centering has been performed per column. All the different DCE categories have been tested. Symbols inside cells demonstrate the significance level of the outcome of each test (*:0.05; **:0.01; ***:0.001). Significance has been assessed by a non-parametric, permutation-based test.

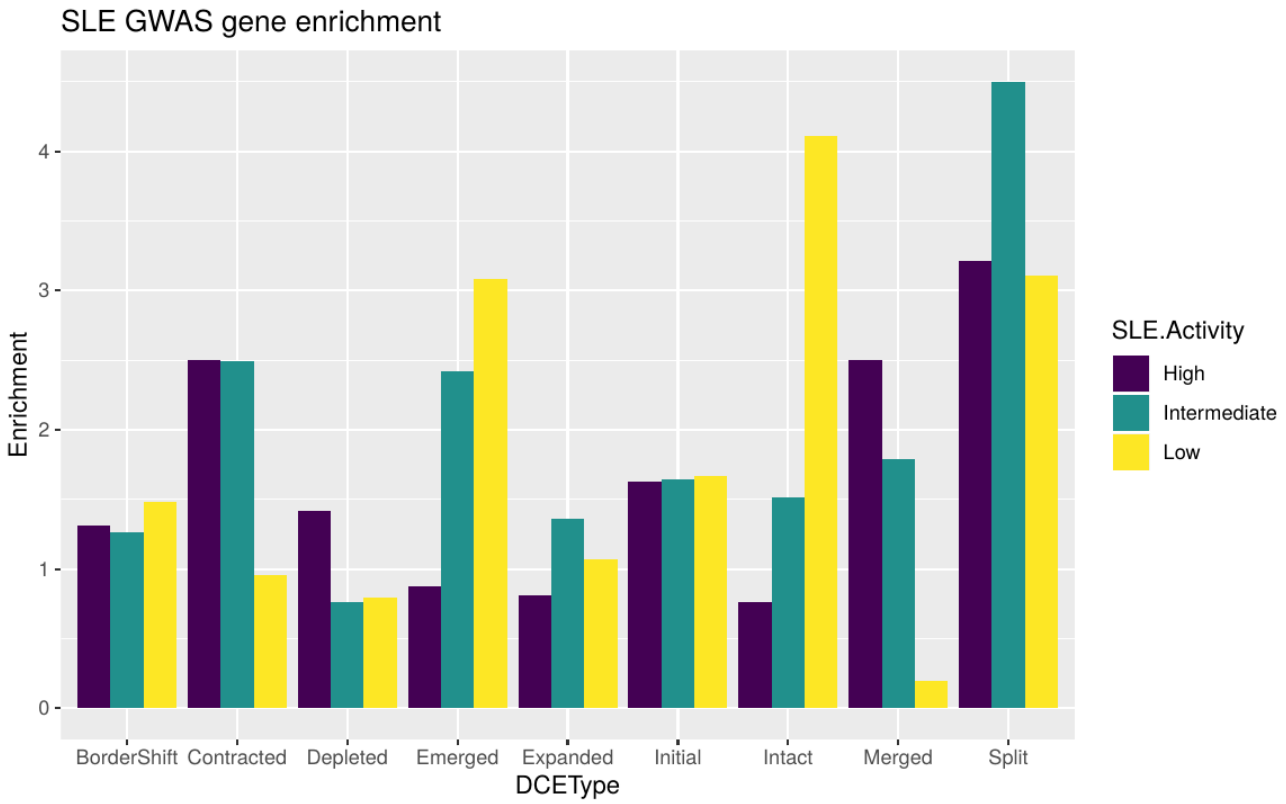


**Supplementary Figure 6. Enrichment of SLE genetically associated genes among different DCE types.** Bars correspond to fold-enrichment of overlap between a set of genes reported as genetically associated with SLE (downloaded from GWAS Central, December 2019) against different types of DCE for each of the three disease activity groups. All values >2 were significant at p-value<0.01.

### Supplementary Methods

#### Analysis of Gene Expression

We obtained RNA sequencing data from a total of 142 SLE patients and 58 healthy individuals originally published in (1). Both groups contained individuals mainly of Caucasian ethnicity and an approximate ratio of 1:5 for male over female. The sequencing material was derived from whole blood samples. Moreover, it should be noted that patients suspended lupus medication 12 hours preceding sampling. Extensive information regarding patient characteristics, mRNA extraction, sequencing protocol, quality control and mapping are thoroughly reported by (1).

We used FeatureCounts (2) to extract raw counts and quantify expression levels for a comprehensive set of human genes, as compiled under the GENCODE annotation v15 (https://www.gencodegenes.org/human/release_15.html, GRCh37). A fragment was counted in case of any overlap with an exon feature and the counts were grouped based on the "gene_name" attribute of the annotation entities. Only fragments with both ends successfully mapped were considered for summarization. Fragments that were chimeric, overlapping multiple meta-features (genes), not uniquely mapped, or having any read marked as duplicate were discarded.

The initial number of genes included in the raw count table was 51716. A multi-step filtering approach was adopted. At first, the "type" of each gene was extracted from the annotation GTF file used in fragment summarization. Then, genes belonging to any of the following types were filtered out: "pseudogene", "processed transcript", "polymorphic pseudogene", "antisense", "sense intronic", "sense overlapping", IG_V pseudogene", "IG_C pseudogene", "TR_V pseudogene", "TR_J pseudogene", "IG_J pseudogene", "non_coding", "Mt-tRNA" and "Mt-rRNA". The total number of genes belonging to those categories were 20190. Subsequently, 167 genes, which had multiple entries in the annotation file, with the same "gene_name", but different chromosome attribute, and could therefore generate errors in the fragment summarization process, were removed from our dataset as well. The number of genes that passed the filtering procedure was 31318.

At the final stage, a two step normalization was implemented on raw counts (filtered for the different irrelevant gene types), using relative log expression (RLE) followed by normalization for gene length.

#### Stratification of the patient cohort

We grouped patient samples according to a clinical SLE disease activity index (SLEDAI) (3). A value of 0 for SLEDAI indicates inactive patients and it increases with higher disease severity. Thus, we used it to assess groups of diverse disease activity. Patient samples were separated into three groups. A low disease activity group, with a maximum SLEDAI value of 2, an intermediate, with SLEDAI indices that ranged from 3 to 8, and a high disease activity group with

SLEDAI indices greater than 8. The number of samples in each group were 55, 61 and 26 respectively. Both Differential and Topological analyses of gene expression have been performed on these three groups in comparison to the healthy group.

**Differential gene expression analysis**

Differentially expressed genes (DEGs) were called using zero inflated generalized linear models provided by MDSeq tool (4). For this analysis we applied an additional filtering layer. Genes with a mean cpm value lower than 0.05 were excluded. The remaining genes were 18447. Furthermore, we incorporated gender and drug treatment as covariates in our models. DEGs were identified based on both statistical significance and effect size. They were defined as genes with corrected p-value lower than or equal to 0.05 and absolute log2(Fold-Change) value greater than or equal to 0.5.

**Modular Analysis of Differential Gene Expression**

We followed a gene set enrichment approach, in order to investigate the over-representation of specific functional modules in our dataset. In a gene set enrichment analysis, the objective is to detect functional modules, whose gene members tend to cluster towards the top (or bottom) of a ranked list. Here, we ranked genes according to absolute differential expression values ($\log_2$|Fold-Change|), and we tested the over-representation of functional groups of genes (modules). The tested modules are closely related to blood tissue and immunity, as identified by two independent studies (9, 10), that analyzed a plethora of blood gene expression datasets in a variety of conditions. Finally, we assessed statistical significance by applying a CERNO test (8). We filtered modules according to statistical significance (corrected p-value ≤ 0.05) and their DEG content, i.e. at least 15% of gene members had to be DEGs according to our previous analysis.

**Weighted Gene Co-expression Network Analysis (WGCNA)**

In order to detect modules of genes with correlated expression independently of their genome topology, we implemented weighted gene co-expression network analysis (WGCNA) (5). Briefly, WGCNA represents genes as nodes in a network. These are connected to each other by edges, to which an adjacency score is attributed. The adjacency score of a node pair is calculated by a power function of the absolute value of the correlation of the corresponding pair of genes. Here, the soft threshold parameter (the power in the adjacency function) was selected to be 10 according to scale free topology criterion, which suggests choosing the lowest possible value such that approximate scale free topology is reached in the network. Modules of co-expressed genes were extracted from the network based on the hierarchical clustering of topological overlap measure and the subsequent implementation of a dynamic cutter. Those initial modules were merged using hierarchical clustering of their eigengene vectors and by cutting the resulted tree at the height of 0.25. The final modules were functionally characterized by utilizing pathway enrichment and calculating the correlations of the module eigengene vectors with a variety of clinical traits.

**Robust co-expression matrix calculations**

Each chromosome was split in 10kb bins, starting from the start of the first gene, till the end of the last gene. Using the normalized counts of genes attributed to a bin, the mean count was calculated for each individual belonging to the same sample group (healthy, low, intermediate and high SLE activity). We then calculated the Spearman correlation matrix between all bins that resided in the same chromosome for all samples within each group. Chromosomal bins with zero expression were ignored for the rest of the analysis. This procedure produced a square correlation matrix for each chromosome. To statistically evaluate the correlation coefficients, a Monte-Carlo-like approach was implemented. The bin counts, of each individual separately, were shuffled randomly and afterwards the correlation matrix was re-constructed. That procedure was repeated 1000 times for each chromosome. In every iteration the calculated correlation coefficients were compared to the original correlation coefficients that were calculated using the intact bin counts. The p-value for each coefficient was set as equal to the fraction of those 1000 permutations, in which the corresponding coefficient had the same or more extreme value compared to the actual one. The correlation coefficients with p-value greater than 0.05 were discarded from the analysis (turned into 0s).

**Cell type estimation and entropy calculation**

We used the results of CIBERSORT (7) for the estimation of the proportion of different immune cell types in whole blood, as performed in (1) . Shannon entropy was used as a metric, in order to assess the variability/uncertainty in the proportions of the different cells types between healthy and SLE subjects.

$$H=-\sum_{i=1}^{n} p\left(x_i\right)\cdot \log_2\left(p\left(x_i\right)\right)$$

where $H$ is the Shannon (information) entropy, $p(xi)$ is the estimated proportion of $x_i$ cell type in whole blood and $n$ is the total number of estimated cell types. Entropy was calculated for every individual in the dataset. Subsequently, the difference between the distribution of entropies of healthy and SLE groups were statistically evaluated by a non parametric Wilcoxon–Mann–Whitney test.

**DCE Analysis**

Two different metrics were applied to explore the differences of DCE sets of different groups. DCEs were handled as a set of chromosomal intervals. The first metric used was the Jaccard similarity coefficient. DCE pairs between two different groups (e.g. healthy and patient groups) with chromosomal coordinates that overlap were detected. For every pair the Jaccard index was calculated. The second metric used was the BP distance score (6). BPscore takes into account the relative chromosome size and thus may provide a more nuanced assessment of coordinate similarity. For the calculation of BP-score we used a publicly available python script, (https://github.com/rz6/bp-metric), that is provided by the authors.

**"Disruptor" definition**

"Disruptor" genes are defined as those, which reside between two "patient DCEs" derived from a split of a "healthy DCE". More precisely, for each "split" characterized healthy DCE, attributed genes were listed. Subsequently, they were compared as a set with those genes, attributed to patient DCEs, corresponding (overlapping) to the initial healthy DCE. Genes absent from the patient DCEs and present in the healthy DCE were characterized as disruptors. Moreover, those genes were filtered, in order to capture only the genes located in the area between the patient DCEs (the locus of the "split" event). That process was repeated for every distinct patient group.
